# Pseudomonas Putida Dynamics of Adaptation under Prolonged Resource Exhaustion

**DOI:** 10.1101/2023.07.31.551210

**Authors:** Jonathan Gross, Sophia Katz, Ruth Hershberg

## Abstract

Many non-sporulating Bacterial species survive prolonged resource exhaustion, by entering a state termed long-term stationary phase (LTSP). Here, we performed LTSP evolutionary experiments on the bacterium *Pseudomonas putida*, followed by whole genome sequencing of evolved clones. We show that *P. putida* is able to persist and adapt genetically under LTSP. We observed a gradual accumulation of mutations within the evolving *P. putida* populations. Within each population, independently evolving lineages are established early on and persist throughout the four-month-long experiment. Mutations accumulate in a highly convergent manner, with similar loci being mutated across independently evolving populations. Across populations, mutators emerge, that due to mutations within mismatch repair genes developed a much higher rate of mutation than other clones with which they co-existed within their respective populations. While these general dynamics of the adaptive process are quite similar to those we previously observed in the model bacterium *Escherichia coli*, the specific loci that are involved in adaptation only partially overlap between *P. putida* and *E. coli*.

**Significance statement:** Bacteria often face conditions of prolonged nutrient limitation, following periods of growth. One strategy for dealing with this challenge is entry into a state termed long-term stationary phase (LTSP), in which a small minority of cells within a population can survive and persist, by recycling the remains of their deceased brethren. Here, we broaden our understanding of adaptation under LTSP, by studying it in the bacterium *Pseudomonas putida*. We show that many of the dynamics of LTSP genetic adaptation are quite general, as reflected by great similarity to what was previously observed in the model bacterium *Escherichia coli*. However, the specific loci that are involved in adaptation substantially vary between *P. putida* and *E. coli*.

## Introduction

Bacteria must often face long periods of severe nutrient limitation, following periods of growth. While some species deal with this challenge by generating spores, most bacterial species cannot sporulate, and some face this challenge by entering a state of growth termed long-term Stationary Phase (LTSP) (Zambrano, et al. 1993; Finkel 2006; Avrani, et al. 2017). Contrary to the impression given by the word ‘stationary’, bacterial populations do not remain stagnant during LTSP. Instead, they continuously undergo physiological, metabolic and genetic changes, even years into their time under resource exhaustion (Finkel 2006; Avrani, et al. 2017; Katz, et al. 2021; Ratib, et al. 2021; Katz, et al. 2023). Although the ability to enter LTSP is not limited to a single bacterial species (Lostroh and Voyles 2010; Bruno and Freitag 2011; Reichert, et al. 2013; Djebbi-Simmons, et al. 2019; Shoemaker, et al. 2021), the majority of studies on LTSP have been performed on the model bacterium *Escherichia coli* (Zambrano and Kolter 1993; Zambrano, et al. 1993; Finkel and Kolter 1999; Farrell and Finkel 2003; Finkel 2006; Kram and Finkel 2014, 2015; Avrani, et al. 2017; Avrani, et al. 2020; Gross, et al. 2020; Nandy, et al. 2020; Katz, et al. 2021; Ratib, et al. 2021; Katz, et al. 2023)

When *E. coli* is placed into rich media, a short period of exponential growth is followed by a period of decline (often referred to as death phase), after which bacteria enter LTSP. The dynamics of *E. coli* LTSP adaptation were explored in a recent evolutionary experiment, in which five populations of *E. coli* were established in July 2015, allowed to enter LTSP and have been maintained under LTSP ever since (Avrani, et al. 2017; Katz, et al. 2021). Analyses of sequencing data from hundreds of clones, sampled from these populations, over nine time points spanning their first three years spent under LTSP revealed several interesting aspects of the dynamics of *E. coli* LTSP adaptation: It was shown that *E. coli* undergoes continuous genetic adaptation under LTSP, through the gradual accumulation of mutations. In three of the five populations mutators emerged. These mutators acquired mutations to a mismatch repair gene, leading to an elevation in their mutation rates. Such mutators tended to co-exist within their populations with non-mutator clones, leading to long-lasting variation in mutation rates within single populations. It was further shown that *E. coli* LTSP adaptation tends to occur through soft, rather than hard sweeps, with several genotypes competing for dominance across sampled timepoints (Avrani, et al. 2017; Katz, et al. 2021). LTSP populations were shown to maintain high levels of standing genetic variation, even as they rapidly and convergently adapted to the extreme conditions to which they were exposed. Such maintenance of variation was shown to enable more rapid re-adaptation to previously met conditions (Avrani, et al. 2020; Katz, et al. 2023). Finally, it was demonstrated that Adaptation under LTSP is characterized by high convergence, meaning that the majority of mutations occurring within non-mutator clones, occur within loci that are also mutated in additional independently evolving populations (Avrani, et al. 2017; Gross, et al. 2020; Katz, et al. 2021). Even more strikingly, often the exact same mutations rise to high frequencies across independently evolving populations (Katz, et al. 2021).

The most striking example of convergent adaptation observed in *E. coli* under LTSP involved mutations to the *rpoB* and *rpoC* genes that, together with the *rpoA* gene, encode the RNA polymerase core enzyme. Over 90% of *E. coli* LTSP clones carry a mutation within either *rpoB* or *rpoC* (Avrani, et al. 2017; Katz, et al. 2021). Often the exact same mutations are seen across independently evolving LTSP populations. Intriguingly, when we carried out the same experiments, on the same strain of *E. coli*, but varied a single experimental parameter – growth volume, we found that different specific *rpoB* and *rpoC* mutations arise in a convergent manner, across populations (Gross, et al. 2020). *rpoB* and *rpoC* were found to be targets for adaptation in many additional evolutionary experiments that explored *E. coli* adaptation to a large variety of selective pressures (Conrad, et al. 2010; Tenaillon, et al. 2012; Graves, et al. 2015; Bruckbauer, et al. 2019; Cohen and Hershberg 2022) The specific sites of *RpoB* and *RpoC* that were mutated varied between conditions, with very little overlap found across conditions (Cohen and Hershberg 2022).

*P. putida* is a gram-negative, rod-shaped bacterium belonging to the gamma-proteobacteria class and is widely distributed across various environments (Clarke 1982). It displays a diverse range of metabolic activities, indicating its adaptation to various niches, including the ability to survive in soils and sediments with high levels of heavy metals and organic contaminants (Wackett and Gibson 1988; Zylstra, et al. 1988; Eaton 1997; Choi, et al. 2003). Some strains of *P. putida* are well known for their ability to break down a wide range of natural and artificial chemicals, including benzene, toluene, ethylbenzene, and p-cymene (Nakazawa 2002; Nelson, et al. 2002; Timmis 2002; Dos Santos, et al. 2004). *P. putida* strains also serve as rhizospheric and endophytic bacteria, promoting plant growth (Taghavi, et al. 2009). To date, the genomes of several *P. putida* species have been fully sequenced (Nelson, et al. 2002). The *P. putida* F1 strain, used in this study, is considered a reference strain for the degradation and growth of aromatic hydrocarbons. Moreover, it is a well-established model for investigating bacterial growth under controlled laboratory conditions.

In this study, we carried out LTSP evolutionary experiments on *P. putida* F1, followed by whole genome sequencing of evolved clones. This allowed us to reveal that many of the general dynamics of adaptation of *P. putida* to LTSP resemble those of *E. coli*, suggesting that these dynamics may be quite general. Specifically, mutation accumulation rates within non-mutators were similar, the patterns of emergence of mutators was similar, there where high levels of convergence and adaptation occurred via soft, rather than hard sweeps, leading to the establishment and co-existence of lineages within individual populations. At the same time, there was only partial overlap in the specific loci involved in adaptation to LTSP, between the two bacterial species.

## Results

### Pseudomonas putida survives and adapts genetically under prolonged resource exhaustion

To examine whether and how *P. putida* survives and adapts under prolonged resource exhaustion, we performed LTSP evolutionary experiments, following a similar experimental setup to that previously used in our *E. coli* LTSP evolutionary experiments (Avrani, et al. 2017; Katz, et al. 2021). Specifically, we established three independent populations of the model bacterium *P. putida* F1. Each population was initiated by inoculating ∼ 10^6^ cells per ml into a total volume of 400 ml of fresh Luria Broth (LB) inside two-liter Erlenmeyer flasks, which were placed into an incubator-shaker at 225 rpm. The incubator was set to the *P. putida* permissive growth temperature of 30°C. We sampled each population across all experiments, first daily, then weekly, and eventually monthly, up to day 288. We did not add new resources to the cultures throughout the experiment, besides distilled water to counteract evaporation. We quantified the number of viable cells at each sampling point by plating appropriate dilutions onto LB plates and counting the resulting colonies. As a reference, we also plotted in **Figure 1** the *E. coli* measurements, which were gathered as part of our previous work (Katz, et al. 2021). The remainder of each sample was archived at -80°C for future sequencing and additional analyses.

**Figure 1.**
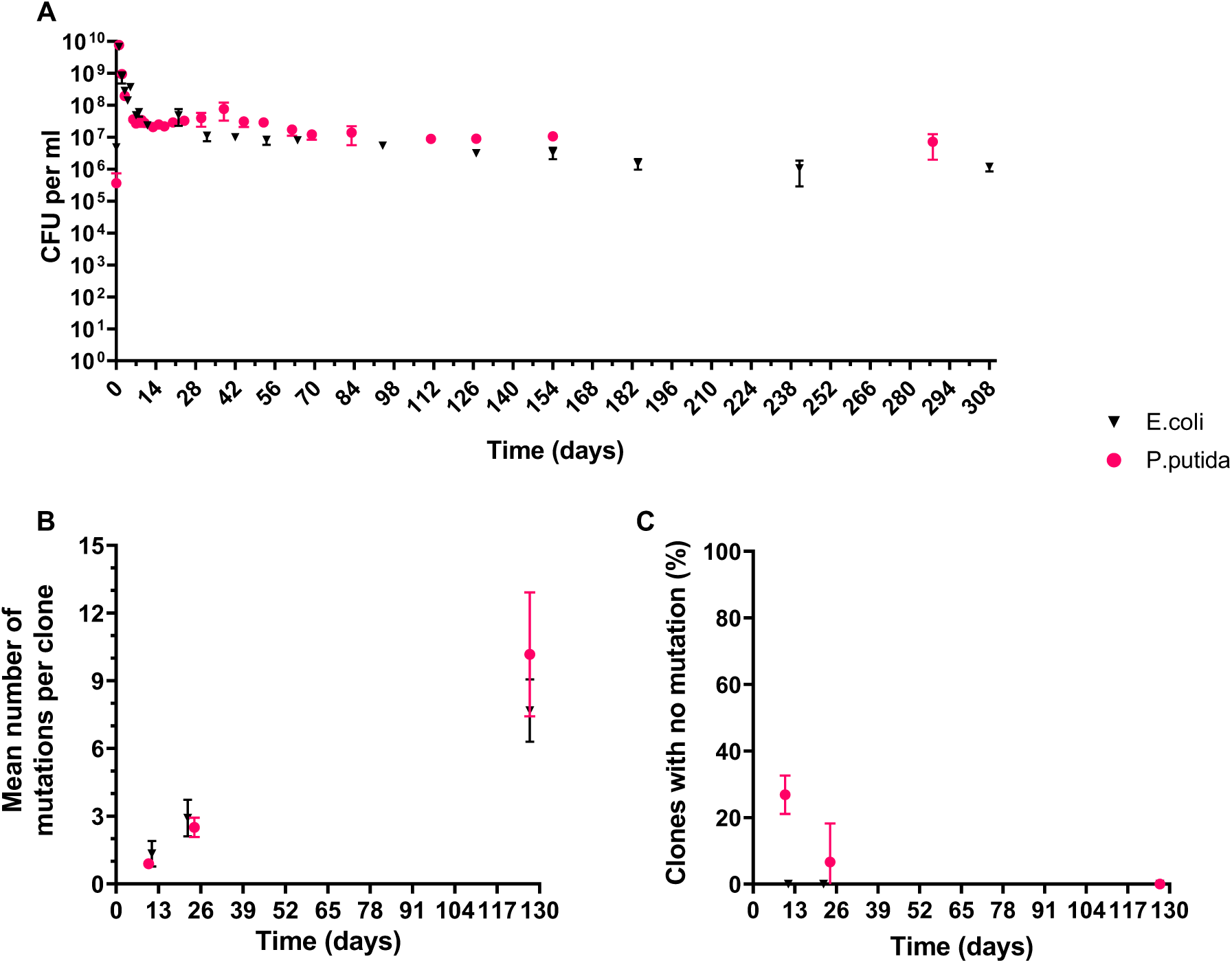
Viability and mutation accumulation under LTSP in *P. putida* (red dots), and *E. coli* (black triangles). (A) Mean viability (cells per ml), as estimated through quantifying colony forming units (CFUs). (B) Mean mutation numbers per sequenced clones, as a function of time. (C) Mean number of clones maintaining an entirely ancestral genotype, across the three populations. In all three plots, error bars represent the standard deviation around the means.

As shown in **Figure 1A**, the growth curve of *P. putida* is quite similar to that of *E. coli*. The average number of *P. putida* cells, across the three populations, grows during the first 24 hours of growth from ∼3.7*10^5^ cells per ml, to ∼7.7*10^9^ cells per ml. *P. putida* populations then enter a rapid death phase, reflected in sharp reductions in the number of viable cells. The rate of this reduction diminishes dramatically, starting at day 10, which we consider to be the first day of LTSP.

Next, for each of the three *P. pudia* populations, we fully sequenced 9-12 clones per population at each of three time points: days 10, 24, and 127. Altogether, we sequenced 95 evolved clones (**Table S1**). The ancestral clones from which each population was established were also sequenced, allowing us to identify the mutations that arose during our experiments. A complete list of all identified mutations can be found in **Table S2**.

In all three *P. pudia* populations, we observed the emergence of mutators, with mutations to the mismatch repair gene, *mutL*. The mutator clones accumulated a higher number of mutations relative to non-mutators. At first, we excluded any mutator clones and calculated the mean number of mutations accumulated by all other clones, as a function of time spent under LTSP. As can be seen in **Figure 1B and Figure S1**, mutations within non-mutator clones accumulate gradually with time, at rates that are quite similar to those previously observed for *E. coli*. An alternative method to assess the accumulation of mutations is to calculate the proportion of sequenced clones that have yet to acquire any mutations and thus maintain their ancestral genotype. Ancestral genotype clones were maintained at the first two time points. No such clones were seen by day 127 **(Figure 1C)**.

Two lines of evidence suggest that many of the mutations accumulated under LTSP in non-mutator clones are adaptive. The first line of evidence is the great convergence with which these mutations accumulate across independently evolving populations **(Figure 2A)**. Across populations, a majority of mutations found within non-mutator clones, occur within loci that are mutated in at least two of the three populations. The second line of evidence for the adaptive nature of many of the mutations observed within our LTSP clones is the enrichment in non-synonymous vs. synonymous mutations relative random expectations. This enrichment is reflected in the ratio of the rates of nonsynonymous to synonymous mutations (dN/dS), being significantly higher than 1 across populations **(Table 1)**.

**Figure 2.**
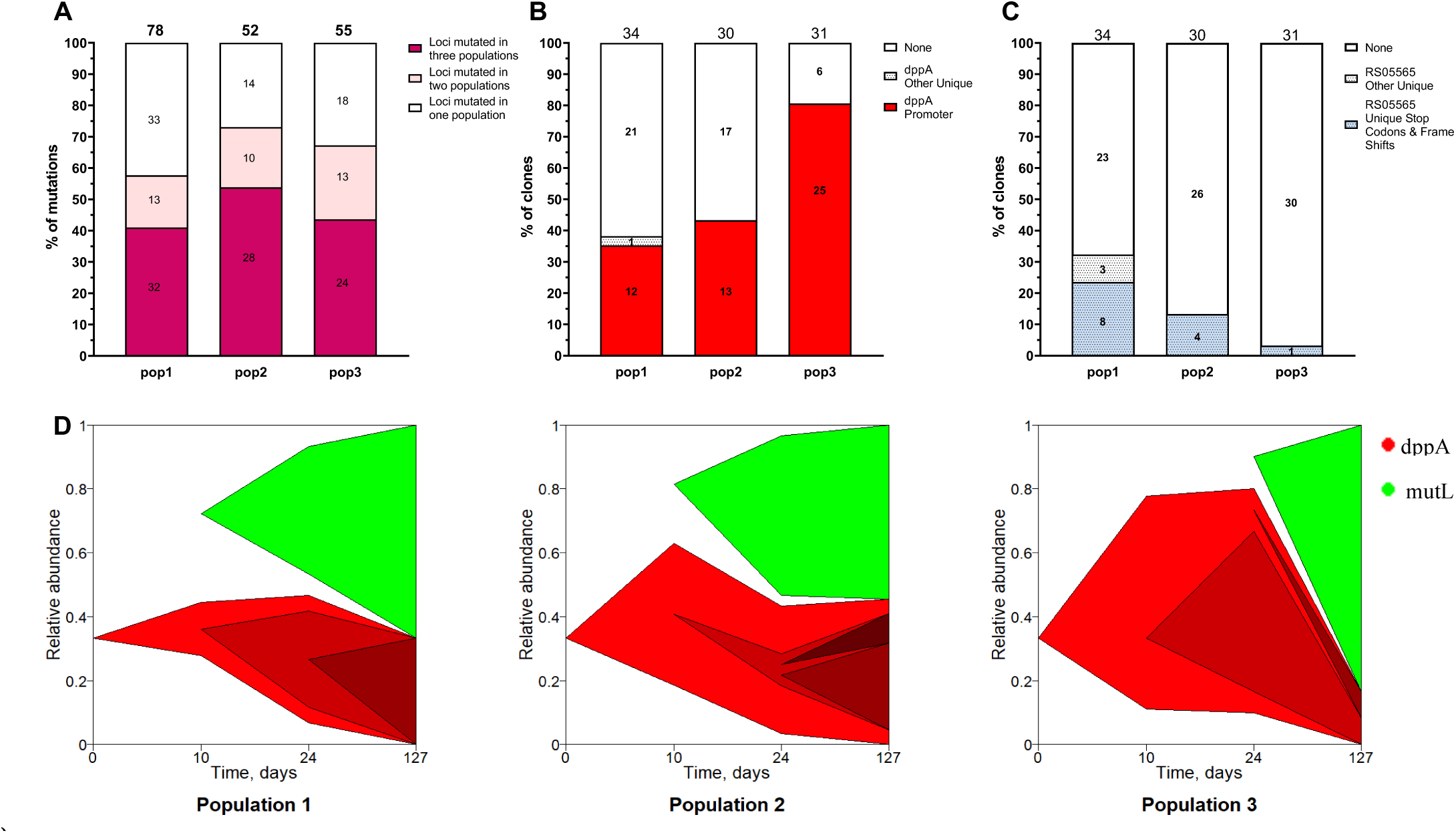
*P. putida* LTSP adaptation is characterized by high convergence and the establishment of lineages. (A) The majority of mutations occurring within *P. putida* non-mutator clones occur within genes that are mutated in a convergent manner. For each population, the number of mutations falling within genes mutated in one, two, or all three populations is presented. Numbers above bars indicate the total number of mutations identified for non-mutators within the population represented by that bar.(B) The *dppA* gene serves as an illustrative example for loci that exhibit specific convergent mutations, likely indicating a specific change in function. Supplementary figure S2 presents all four best examples of such loci. (C) *PPUT RS05565* is an illustrative example of genes that exhibit convergent deactivation across populations. Supplementary figure S3 presents all three examples of such genes. (D) The establishment of lineages and a clear pattern of their co-existence can be observed across all three populations under LTSP. Across all populations two dominant lineages were identified. Muller diagrams, which depict the relative frequencies of these different lineages within LTSP populations 1, 2, and 3, were generated to illustrate the observed patterns. The x-axis represents the sampling times, with mutations that appeared in 30% or more of the population’s clones during at least one time point being used to generate each plot. Variations occurring within the mutator clones were not depicted due to their extensive nature. Muller plots were produced using the R package MullerPlot (Farahpour, et al. 2016).

**Table 1.**
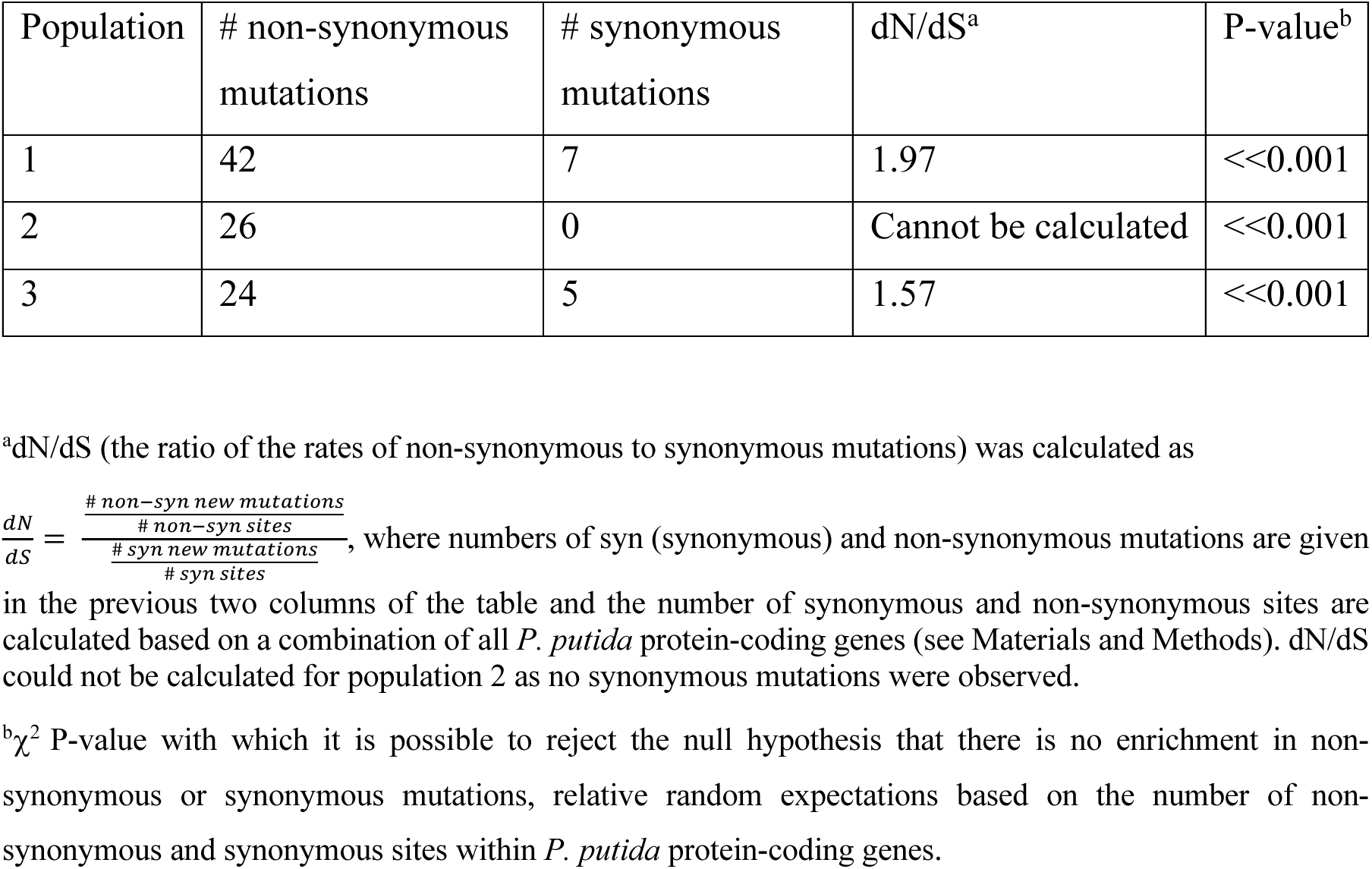
Enrichment in non-synonymous, vs synonymous mutations relative random expectations, for non-mutator clones, across all populations.

| Population | # non-synonymous mutations | # synonymous mutations | dN/dS<sup>a</sup> | P-value<sup>b</sup> |
| --- | --- | --- | --- | --- |
| 1 | 42 | 7 | 1.97 | <<0.001 |
| 2 | 26 | 0 | Cannot be calculated | <<0.001 |
| 3 | 24 | 5 | 1.57 | <<0.001 |

<sup>a</sup>dN/dS (the ratio of the rates of non-synonymous to synonymous mutations) was calculated as
in the previous two columns of the table and the number of synonymous and non-synonymous sites are calculated based on a combination of all *P. putida* protein-coding genes (see Materials and Methods). dN/dS could not be calculated for population 2 as no synonymous mutations were observed.
<sup>b</sup> $\chi^2$ P-value with which it is possible to reject the null hypothesis that there is no enrichment in non-synonymous or synonymous mutations, relative random expectations based on the number of non-synonymous and synonymous sites within *P. putida* protein-coding genes.

### Convergently mutated loci are enriched for functions related to transport and cell wall organization

Utilizing *P. putida* gene annotations found in BioCyc Database (Karp, et al. 2019), we found that genes that are convergently mutated in two or more of our populations are enriched for functions related to cell wall organization and transport. While only ∼12% of *P. putida* genes are annotated as being involved in such functions, 44.3% of the genes we found to be convergently mutated under LTSP are so annotated. This constitutes a statistically significant enrichment (*P* << 0.001, according to a *X*^2^ test).

Based on the types of mutations we observed within convergently mutated loci, we could classify some of them according to whether they likely undergo an adaptive change in function or an adaptive loss of function. Loci for which the same exact mutations are observed across independently evolving populations likely underwent a specific adaptive change of function. At the same time genes in which non-specific deactivation (nonsense or frameshift) mutations are seen across different populations likely experienced an adaptive loss of function (Katz, et al. 2021). Illustrations of the most striking examples of each of the two types of loci are provided in **Figure 2B** and **Figure 2C** respectively. Additional examples are provided in **Figure S2** and **Figure S3**.

The most striking example of an apparent adaptive specific convergent change in function involves the RS04685 gene. This gene is an ortholog of the *E. coli dppA* gene, which encodes a periplasmic binding protein that functions as Part of the ABC transporter DppABCDF involved in dipeptide transport. 54% of all clones sequenced in our analysis carry a mutation in RS04685 or its promoter region, and the same exact mutation within the promoter region of this gene is seen in ∼98% of these clones, across all three populations (**Figure 2B**). Interestingly, in four of the clones that carry this specific RS04685 promoter mutation, extracted from two different populations, we also saw between one and seven additional mutations to the same gene that fall between positions 338 and 350 of its protein sequence. Two of these clones, extracted from two independently evolving populations, shared between them six such mutations and over all four clones, a total of eight different mutations were observed. Six of these eight mutations were synonymous and two were non-synonymous (**Table S2**), suggesting a possible effect at the level of the transcript rather than at the protein level. Notably, *dppA* was also convergently mutated under LTSP in *E. coli* (Katz, et al. 2021), suggesting that this same locus may be important for LTSP adaptation in both organisms.

Additional loci in which the same specific mutations are seen across multiple *P. putida* populations include (1) the promoter region of RS08200, a gene encoding a subunit of the cytochrome c oxidase, which takes part in the aerobic electron transfer chain **(Figure S2B)**; (2) The gene RS04675, a member of the *OprD* family of purine transporters, in which the most frequently observed convergent mutation changes the start codon from TTG to ATG **(Figure S2C)**. Such changes to the start codon were previously shown to affect translation efficiency (Sussman, et al. 1996); and (3) the ABC dipeptide transporter gene RS04670 **(Figure S2D).**

Examples of genes in which deactivation mutations occur in a convergent manner include: (1) The penicillin-binding protein gene, RS05565 **(Figure 2C)**; (2) *RS21800* (**Figure 3SB**) - a cytoplasmic membrane protein that functions as a subunit of RND permease efflux, which is a family of transporters that represent a prominent class of efflux pumps found primarily in Gram-negative bacterial species (Nikaido and Takatsuka 2009); and (3) an ortholog of the *E. coli nrdR* transcription factor (**Figure S3C**), which in *E. coli* is thought to function as a negative regulator of bacterial growth (Torrents, et al. 2007). It is interesting to note that, while *nrdR* does exist in both *P. putida* and *E. coli*, we did not find any mutations within this gene arising under LTSP in *E. coli*.

### Lineage structure of P. putida LTSP populations

We used the genome sequences we obtained to generate trees depicting the relationships between the different clones within each population. This enabled us to generate Muller plots that depict the lineage structure of each population **(Figure 2D)**. These plots show that within each population, lineages are established early and persist throughout the first 127 days of our experiment. Interestingly, each population seems to contain two major lineages that compete with each other. The first of these two lineages is a non-mutator lineage, which is established by clones carrying the same specific mutation, across all populations (The highly frequent, highly convergent mutations within the promoter of the *dppA* ortholog RS04685 gene, as described above). The second lineage is a mutator lineage, in which clones carry a different *mutL* mutation, within each of the populations. In population 3, we saw the highly convergent *dppA* / RS04685 promoter mutation occurring on the background of the mutator lineage as well **(Table S2)**, further highlighting the great convergence with which this mutation tends to occur.

### Mutators dynamics within P. putida LTSP populations

As already mentioned, we observed the emergence of mutator clones in all three LTSP *P. putida* populations. These mutator clones acquired mutations within the mismatch repair gene, *mutL*. In populations 1 and 2, mutator clones were first observed at day 24, whereas in population 3, we first detected mutators only at day 127 **(Figure 3A-C)**. Mutator clones, across all populations, accumulate higher numbers of mutations, compared to non-mutator clones **(Figure 3D-F)**. Population 1 mutator clones acquired a substantially higher number of mutations, than mutators from both other populations. We found that all of the sequenced mutator clones from population 1 carry, in addition to their *mutL* mutation, a mutation within the *dnaQ* gene. The gene *dnaQ* encodes the epsilon subunit of DNA polymerase III, a proofreading exonuclease responsible for correcting errors during DNA replication (Echols, et al. 1983; Scheuermann, et al. 1983). Mutations within *dnaQ* where previously shown to enhance mutation rates within *E. coli* LTSP mutators as well (Katz, et al. 2021).

**Figure 3.**
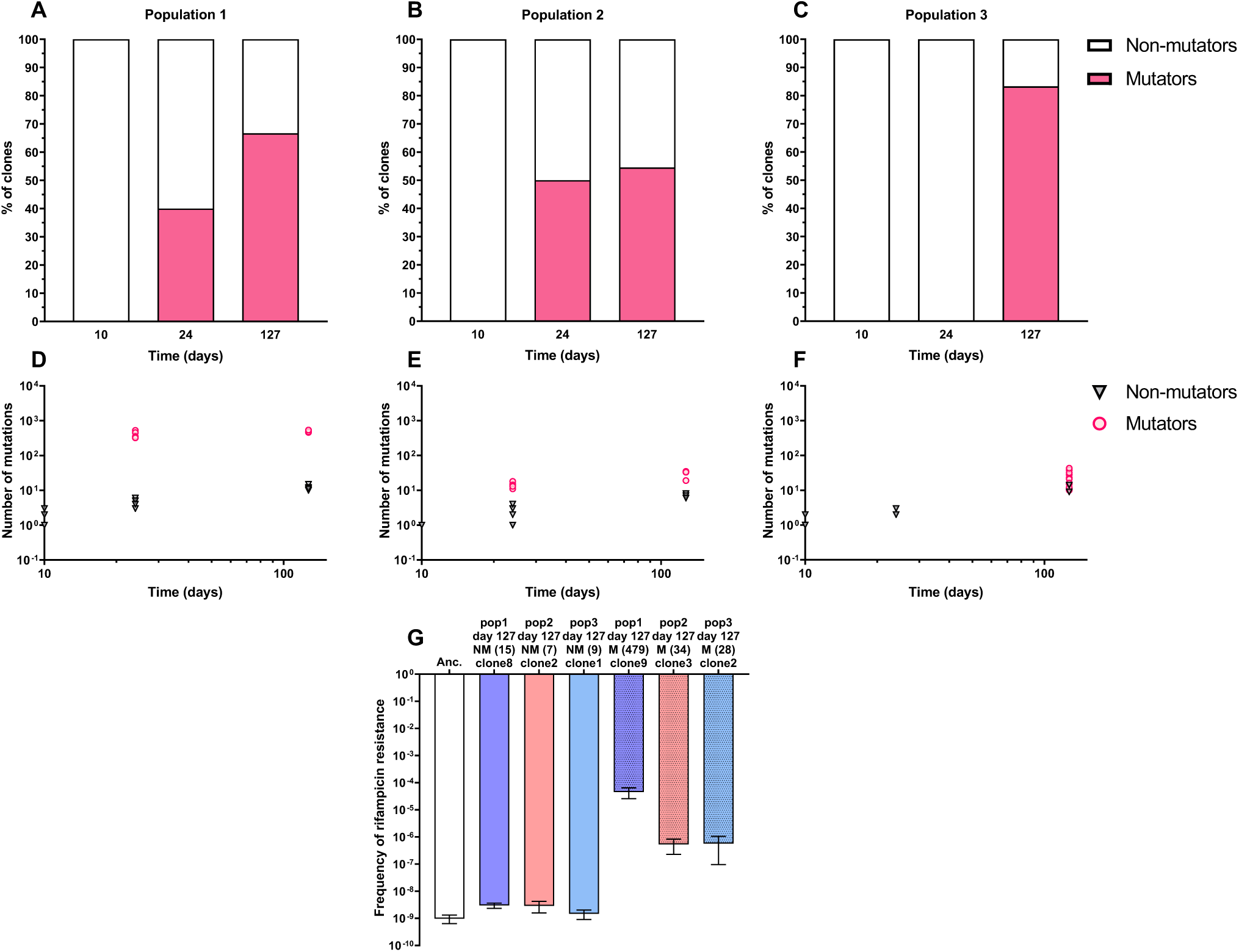
Dynamics of mutator evolution of *P. putida* under LTSP. LTSP adaptation allows the coexistence of mutator and non-mutator clones over extended periods, as evidenced by the observed relative frequencies of mutators in populations 1, 2, and 3 (A-C). The number of mutations accumulated by individual clones over time spent under LTSP in each population. Pink and black dots represent mutators and non-mutators respectively (G). Mean frequencies with which different types of clones develop rifampicin resistance, following overnight growth in rich media without rifampicin. Each mean value was obtained from 15 individual experiments. Error bars represent standard deviations around these means. The number of mutations accumulated by each clone during 127 days under LTSP is indicated in parentheses, NM (Non-mutators), M (mutators), ANC (ancestral *P. putida* strain).

To directly estimate the relative mutation rates of each clone type, we quantified the frequencies at which each type of clone developed resistance to the antibiotic rifampicin after overnight growth in fresh LB without rifampicin **(Figure 3G and Table S3)**. We found that the relative mutation accumulation observed under LTSP, correlates very well with the relative estimated mutation rates. Non-mutator clones from all populations have the lowest mutation rates and lowest mutation accumulation. Mutator clones from populations 2 and 3, have ∼290 times higher mutation rates, and mutator clones from population 1, which also carry a *dnaQ* mutation, have a mutation rate which is ∼1.5*10^4^ times higher than that of non-mutator clones **(Figure 3G and Table S3)**.

Mutator clones with *dnaQ* mutations are present at high frequencies within population 1, at both day 24 and day 127. Yet despite having very high mutation rates, that have led to the accumulation of an average of 413 mutations by day 24, there appears to be very little additional mutation accumulation in the 103 days between day 24 and day 127 **(Figure 3D-3F)**. This lack of mutation accumulation cannot be explained by a lowering of the mutation rates of the mutator clones between days 24 and 127, as mutation rates remain constant between both time points (as estimated by quantifying the frequency of rifampicin resistance, following overnight growth, **Figure 4A**). Comparisons of the mutations present within each of the mutator clones, reveals that clones sampled at day 127 are sometimes much more similar in their mutations to clones sampled at day 24, than to other clones extracted at day 127 **(Figure 4B)**. Combined, this suggests that despite having very high mutation rates, mutator clones persisted within population 1, without accumulating many mutations, between days 24 and 127 (over 100 days in total). A possible explanation for this would be if these clones spent much of their time dormant, between the two time points.

**Figure 4.**
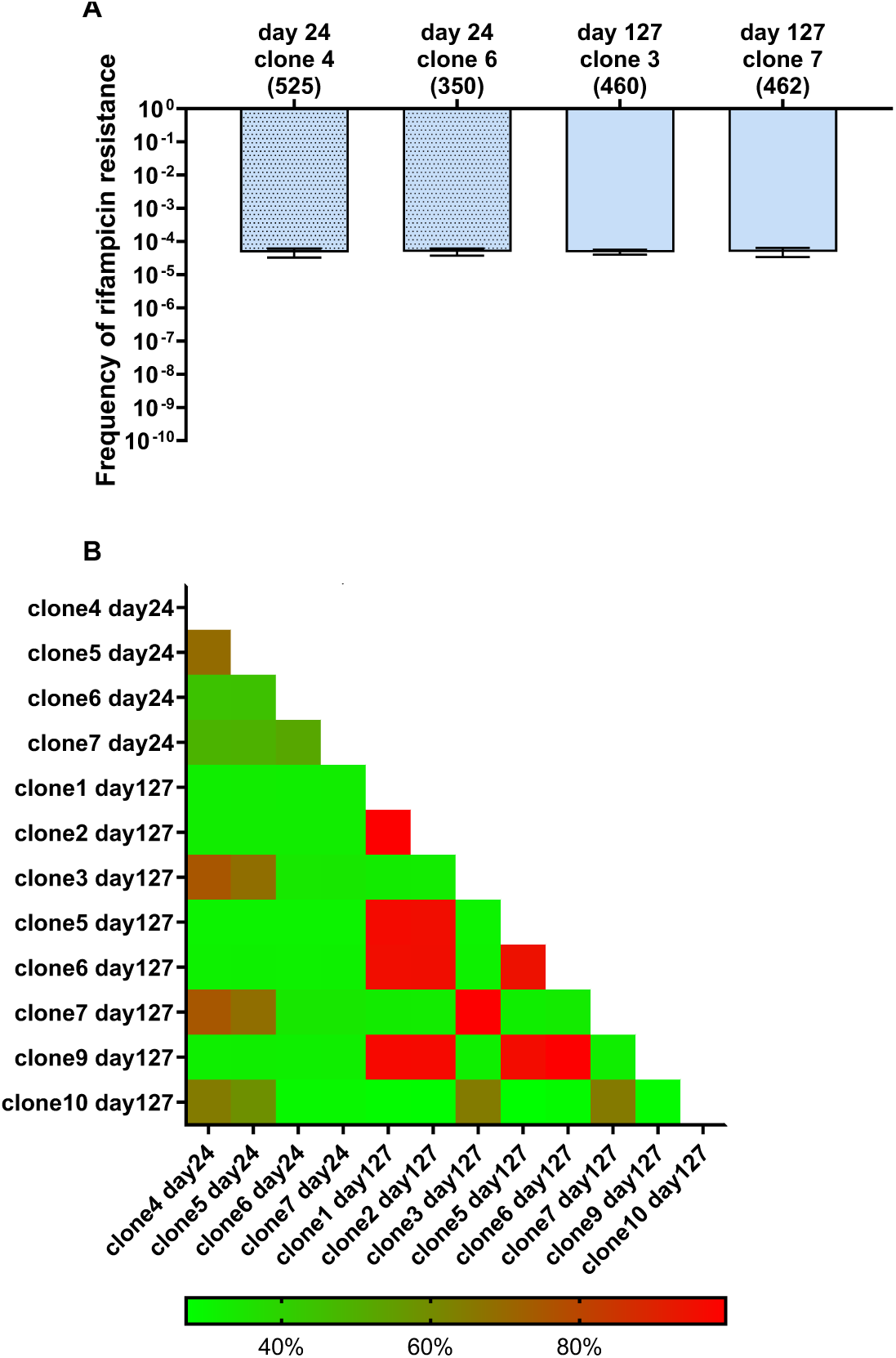
Population 1 mutators maintain high mutation rates throughout the experiment and likely spend some of their time under dormancy. (A) No change observed in mutation rates between mutator clones extracted at days 24 and 127. Presented are the mean frequencies with which different types of clones develop rifampicin resistance, following overnight growth in rich media without rifampicin. Each mean was obtained from 15 individual experiments. Error bars represent standard deviations around the means. The number of mutations accumulated by each clone during 24 or 127 days under LTSP is indicated in parentheses. (B) Day 24 and day 127 population 1 mutator clones are sometimes more similar in their mutation content to each other than to other clones sampled at the same time point. Presented is a heatmap representing the percent of shared mutations between each pair of population 1 mutator clones.

## Discussion

Our results demonstrate that *P. putida* enters, survives and genetically adapts under LTSP, through the gradual accumulation of mutations. Mutation accumulation under LTSP, within non-mutators is dominated by positive selection, as reflected by high convergence and an enrichment in non-synonymous relative synonymous mutations (dN/dS >>1). Many aspects of the dynamics of adaptation under LTSP are general for both *P. putida*, and *E. coli*. In both organisms the growth curve is quite similar, with entry into LTSP occurring at a rather similar timeframe and viability being quite similar, at least during the timeframe studied here (**Figure 1A**). Mutation accumulation rates within non-mutators are also quite similar (**Figure 1B**), although *P. putida* maintains entirely ancestral genotypes within the population, slightly longer than *E. coli* does (**Figure 1C**). In both organisms, mutations occur in a highly convergent manner, across independently evolving populations. The trend by which a majority of mutations occurring within non-mutators fall within loci that are also mutated in additional independently evolving populations is perhaps a little more pronounced in *E. coli*, but nevertheless is clearly true also for *P. putida* (**Figure 2A**).

We observed, in both organisms, the occurrence of mutators, that acquire mutations in mismatch repair genes, resulting in elevated mutation rates and elevated mutation accumulation. In both organisms we observed cases in which the acquisition of an additional mutation within the *dnaQ* gene led to the establishment of mutators with even higher mutation rate. Finally, in both organisms adaptation is characterized by the establishment and persistence of lineages that co-exist within their respective populations, with other lineages, in a pattern of adaptation via soft-, rather than hard-sweeps.

While the general dynamics of genetic adaptation under LTSP were quite similar between *E. coli* and *P. putida*, the identity of the genes that are convergently mutated, and thus likely to be involved in LTSP adaptation, differed substantially between the two species. Genes mutated in a convergent manner in *E. coli* included some of the most central master regulators of gene expression (e.g., the RNA polymerase core enzyme, RNA polymerase’s housekeeping sigma factor RpoD [σ70], and the cAMP-activated global transcriptional regulator CRP) (Katz, et al. 2021). These genes were not mutated in *P. putida*. However, in both organisms convergently mutated genes were enriched for functions related to transport.

The three examples of clear overlap in genes involved in adaptation between the two species were: (1) the mismatch repair genes (*mutL* in the case of *P. putida* and *mutL*, *mutS* and *mutH* in the case of *E. coli*) that lead to the mutator phenotype; (2) *dnaQ* – mutations to which enhance mutator mutation rates even further; and (3) *dppA -* In *E. coli* DppA functions as a periplasmic binding protein for the ABC dipeptide transport system, and its presence is essential for chemotactic behavior (Manson, et al. 1986; Abouhamad, et al. 1991). While *dppA* was shown to be convergently mutated under LTSP in both *E. coli* and *P. putida*, it appears to serve a much more central role in *P. putida* adaptation to LTSP. This is reflected by clones carrying the same exact specific mutation in the promoter of *dppA* being the establishers of the major non-mutator lineage across all three *P. putida* populations.

While the RNA polymerase core enzyme appears to be an important target for LTSP adaptation in *E. coli* (Avrani, et al. 2017; Katz, et al. 2021), no mutations to the RNA polymerase are seen in our *P. putida* LTSP populations. RNA polymerase was shown to serve as a major target for adaptation in *E. coli*, not only under LTSP, but under a very large variety of lab-induced conditions (Cohen and Hershberg 2022). That no such mutations are seen in *P. putida* may suggest that what appears to be a very general trend when looking at the evolution of *E. coli*, may be less general than thought. This highlights the risk in relying heavily on a single model bacterium. It will be interesting to understand why it is that RNA polymerase adaptations seem to be so central, specifically in *E. coli* evolution and to understand in what additional organisms the RNA polymerase constitutes or does not constitute such a hub for adaptation.

Unlike most evolutionary experiments that dilute out slow-growing and dormant cells, LTSP evolutionary experiments allow dormant or nearly dormant cells to persist within their populations. This in turn likely greatly affects the dynamics of adaptation under LTSP. Here, we see a possible example of this in the population 1 mutator clones. Mutator clones within population 1 are first seen at day 24 of our experiment, and are also seen at day 127. Despite carrying both a *mutL* mutation and a *dnaQ* mutation, resulting in mutation rates that are over 1000 times those of non-mutator clones, very little mutation accumulation appears to occur within these clones between day 24 and day 127. Given that there appears to be no difference in the high mutation rates of these clones between day 24 and day 127, a likely explanation for this is if these clones remained largely dormant between the two time points. No similar example of persistent mutator lineages that appear not to accumulate mutations for such long periods of time were observed so far in our *E. coli* LTSP experiments. Additionally, we found that relative to *E. coli, P. putida* populations maintained within them entirely ancestral genotypes for longer periods of time under LTSP (**Figure 1C** and (Gross, et al. 2020)). It is tempting to speculate that these differences between the two species may stem from an increased tendency of *P. putida* to maintain dormancy under LTSP, compared to *E. coli*. However, this possibility will need to be further examined to determine whether this is indeed the case.

## Materials and Methods

### LTSP Evolutionary Experiments

The *Pseudomonas putida* F1 strain used in this study was purchased from the ATCC (strain number 700007). Using this strain and following a similar protocol to that previously described in (Avrani, et al. 2017), three LTSP populations were established. To initiate these populations, individual colonies of *P. putida* F1 were cultured overnight, and three colonies were used to inoculate test tubes containing 4 mL of fresh medium. Each culture was allowed to grow until reaching an optical density (OD) of 1, after which approximately 2 x 10^9^ cells (1 mL) were used to inoculate a 400 mL volume of Luria Broth (LB) in a 2-liter polycarbonate breathing flask. Subsequently, the three flasks were incubated at 30 °C with shaking at 225 rpm. The cultures were not supplemented with any external nutrients or resources over time, except for sterile water, which was added every 10-15 days to compensate for evaporation, based on the weight loss of each flask during the respective time interval.

### Sampling LTSP Populations and Estimating Viability

Initially, one ml samples were collected from each culture on a daily basis, and subsequently on a weekly, and progressively longer basis as depicted in **figure 1A**. Samples were used to prepare dilutions, which were then plated utilizing an automated plater to assess viability through live cell counts. The remainder of the samples were then preserved in a -80 °C freezer by freezing them in 50% glycerol for subsequent analysis.

### Sequencing of LTSP Clones

Thawed frozen cultures of *P.putida* F1 from each population and time point (days 10, 24, and 127) were subjected to dilution plating and overnight growth. To minimize the number of generations each clone undergoes prior to sequencing and reduce the occurrence of mutations during regrowth, 9-12 colonies (as shown in **Table S1**) from each culture were utilized to inoculate 4 mL of medium in a test tube and cultivated until reaching an OD of approximately 1. One milliliter of the culture was subjected to centrifugation at 10,000 x g for 5 minutes, and the resulting pellet was utilized for DNA extraction. The remaining culture was then archived by freezing it in 50% glycerol at -80°C. The Macherey-Nagel NucleoSpin 96 Tissue Kit was employed for DNA extraction, and sequencing library preparation followed the protocol delineated in (Baym et al. 2015). Subsequently, paired-end 150 bp sequencing was conducted utilizing an Illumina HiSeq 2500 machine at the Technion Genome Center. The ancestral clones from which each population was established were also similarly sequenced. The short read data resulting from our sequencing was deposited to the Short Read Archive (SRA), under BioProject ID: PRJNA963207.

### Calling of Mutations

For mutation calling, the reads acquired for each LTSP clone or ancestral strain were mapped to the reference genome of *Pseudomonas putida* F1 (accession NC 009512). The mutation data for an LTSP clone were recorded only if they were found in the LTSP clone’s genome, and not in the ancestral genome. The breseq platform was utilized for sequence alignment and mutation calling. This platform facilitates the identification of various types of mutations, including point mutations, short insertions and deletions, larger deletions, and the formation of new junctions (Deatherage and Barrick 2014).

### Calculating the Proportion of Non-Synonymous Versus Synonymous Mutations Expected Under Neutrality

The DNA sequences of all protein-coding genes in *P. putia* strain F1 were downloaded from the NCBI database. For each position of each gene, we examined the likelihood that a mutation at this position would lead to a non-synonymous or a synonymous change. For example, any change to the third codon position of a 4-fold degenerate codon would be synonymous, so mutations at such a position would be 100% likely to be synonymous. In contrast, mutations to the third codon position of a 2-fold-degenerate codon will be synonymous for a third of possible mutations and non-synonymous for two-thirds of possible mutations. In such a manner, we could add up the likelihood of a mutation being synonymous or non-synonymous across the entirety of *P. putida* F1’s protein-coding genes.

### Quantifying Rifampicin Resistance Frequencies for Individual LTSP Clones Following Overnight Growth in Fresh LB

The method employed to select individual bacterial clones involved the retrieval of samples stored at a temperature of -80°C, followed by streaking of the samples on a solid medium to obtain a single colony. The isolated colony was then cultured in 4 ml of LB overnight, with shaking at 225 rpm and at a temperature of 30°C. Subsequently, 100 µl of the resultant culture was plated on both LB agar plates and LB agar plates supplemented with 100 µg/ml rifampicin (Sigma–Aldrich), with appropriate dilutions. The frequency of rifampicin resistance was determined by dividing the CFUs obtained on the rifampicin-containing plates by the CFUs on the plates that did not have rifampicin added.

## Acknowledgments

This work was supported by an ISF grant (No. 1860/21, awarded to R.H). The described work was carried out in the Rachel & Menachem Mendelovitch Evolutionary Process of Mutation & Natural Selection Research Laboratory.

## Data Availability Statement

Data of all mutations accumulated within the analyzed LTSP populations, at the analyzed time points are provided in **Table S2**. Raw sequencing reads were deposited to the sequence read archive (SRA) BioProject ID: PRJNA963207.

