## Supplementary Figures S1-S3 for "Pseudomonas Putida Dynamics of Adaptation under Prolonged Resource Exhaustion"

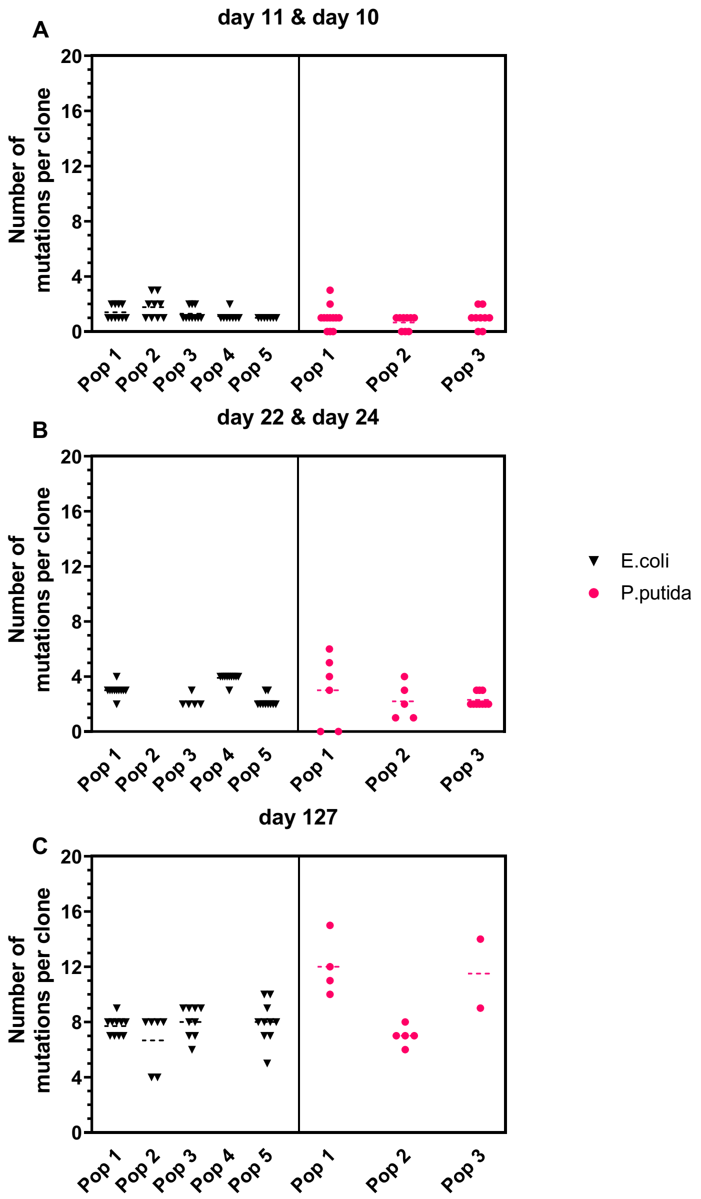


**Figure S1. Number of mutations accumulated per non-mutator clone for *P. putida* clones (pink circles) and *E. coli* clones (black triangles).** Each marking represents an individual clone and each figure bar represents a time point.

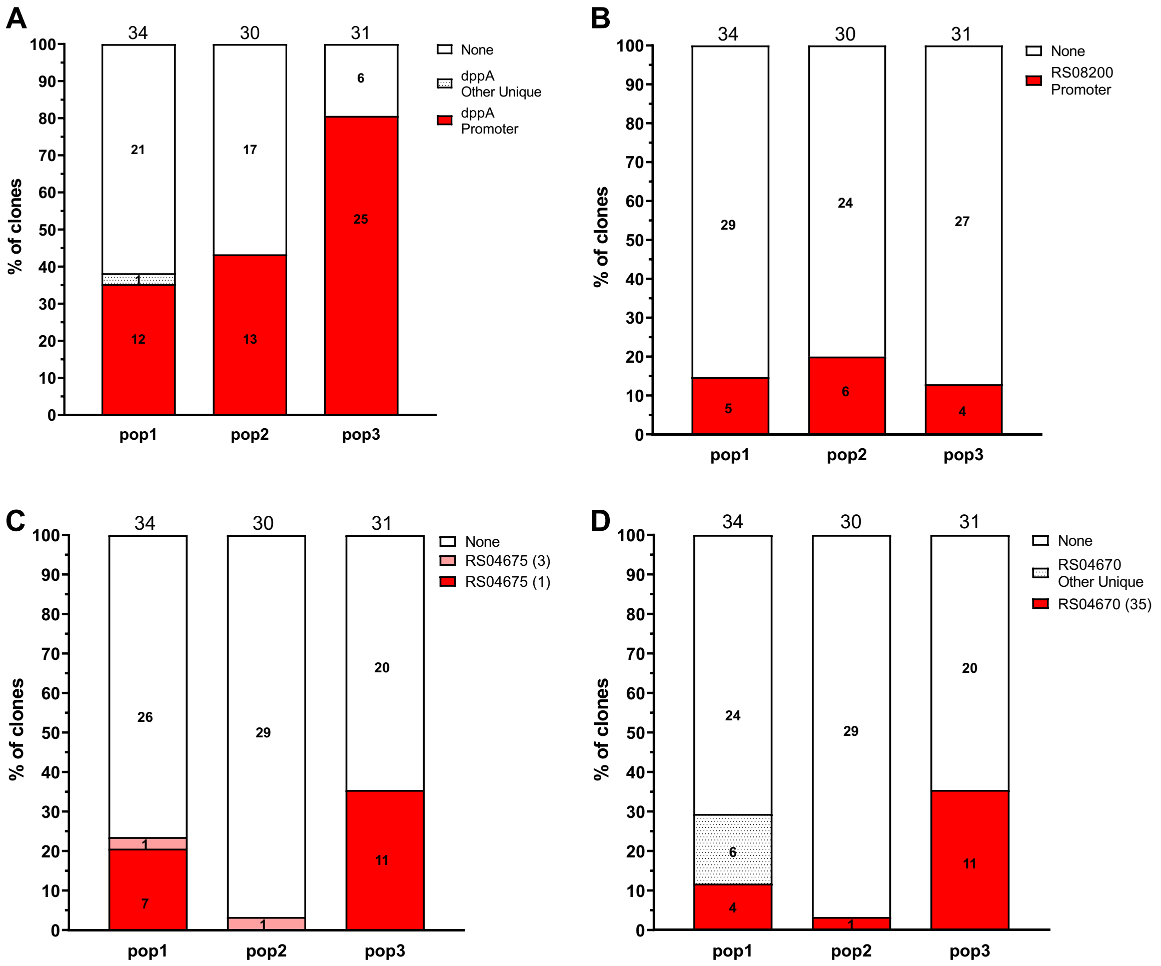


**Figure S2. The loci identified that were found to harbor only one or, at most, two convergent mutations.** (A) *dppA* gene promoter (B) *PPUT RS08200* gene promoter (C) *PPUT RS04675*; (D) *PPUT RS04670*. The positions that contain a mutation in more than one population are assigned a unique identifier, indicated by the protein name and the position number in parentheses. Gene's name contains an asterisk if the mutation fell within the gene's apparent promoter region. Mutations that appear in only one population are clustered under the 'other-unique' category.

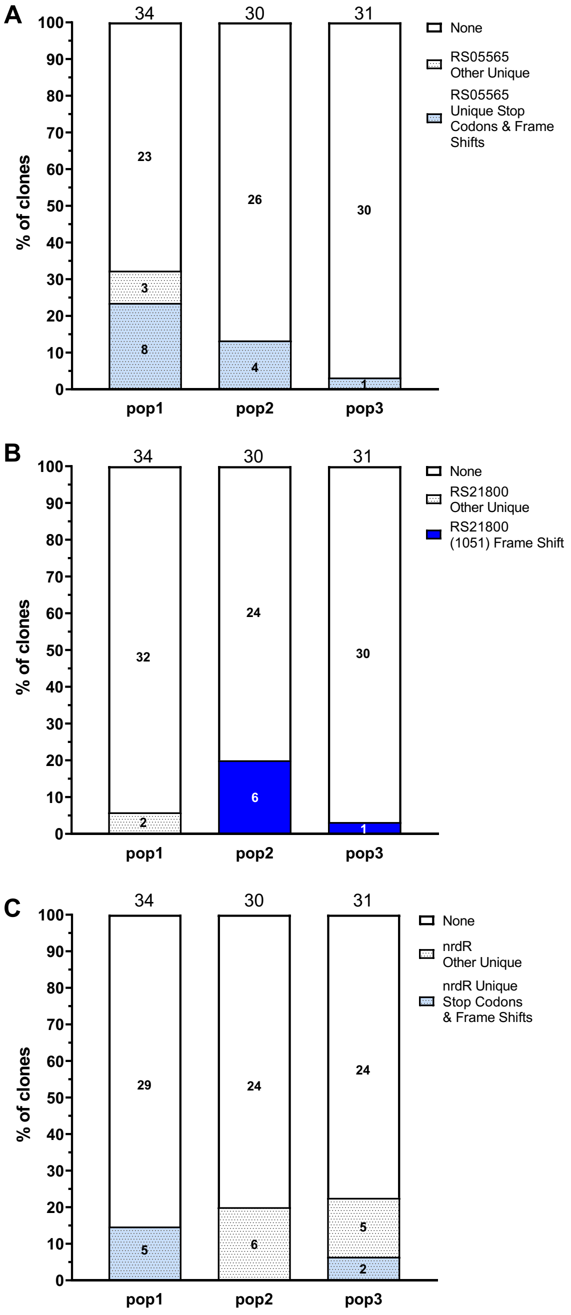


**Figure S3. Genes that are deactivated in a convergent manner.** (A) *PPUT RS05565* (B) *PPUT RS21800* (C) *nrdR*. Positions with mutations that occur in multiple populations are assigned a distinct identifier, which includes the protein name and the position number. If the mutation results in a frame shift or stop codon, it is noted as well. Mutations occurring in only one population are clustered in the 'Unique stop codons & frame shift', or 'other-unique' sections of the population's bar.
